# Optimization of the NFκB-DNA ensemble leads to high affinity binding of partial consensus sites

**DOI:** 10.64898/2026.10.26.762037

**Authors:** Tristan R. Gunther, Sophia M. Hofmeister, Luke Sebastian, Hannah E.R. Baughman, Neville Bethel, Elizabeth A. Komives

## Abstract

Our ability to respond to a myriad of external stressors is dependent on the activity of specific transcription factors that precisely regulate gene expression. While some transcription factors have high specificity for their consensus DNA binding site, others such as NFκB (RelA/p50) readily bind to partial and nonconsensus sites. In fact, ChIP-Seq data shows that nearly half of the DNA-bound RelA occupies sequences that don’t match the consensus. Here we explore how NFκB (RelA/p50) binds to alternative sequences that differ subtly from the consensus. The two subunits of the NFκB heterodimer, RelA and p50, form an extensive pocket where DNA binding is mediated by both base-specific contacts and nonspecific interactions with the DNA backbone. RelA and p50 each form sequence-specific contacts with DNA at different 4-nucleotide sequences separated by two nucleotides, together forming a 10-base pair consensus binding sequence. Partial consensus sequences, known as “half-sites”, match the consensus sequence at only one of these two 4-nucleotide interaction sites. Here, using hydrogen deuterium exchange mass spectrometry (HDX-MS) and molecular dynamics (MD) simulations to investigate protein structural dynamics, we observe that rearranging nucleotides at the RelA-interacting end of a consensus DNA sequence to form a half-site induces dynamic structural changes through both subunits of RelA/p50. MD simulations reveal a shift in the position of DNA within the RelA/p50 binding pocket that allows optimal interaction with the half-site sequence. Our results demonstrate how RelA/p50 fine-tunes its interactions to achieve high-affinity binding with multiple different target DNA sequences.

## Introduction

In response to a multitude of external signals, cells use transcription factors to turn genes on and off in a highly regulated pattern to elicit appropriate responses. Transcription factors recognize and bind to target DNA sequences to activate or repress transcription of their associated genes (1). The Nuclear Factor κB (NFκB) transcription factor family plays a central role in the cellular stress response. Transcription-regulating activity of NFκB is inducible in response to a variety of immune- and inflammation-related stimuli leading to the potent activation of hundreds of genes (2–4). The NFκB family is made up of five proteins that form homo- and heterodimers, but the overall population in most cells is dominated by the RelA/p50 heterodimer (**Figure 1A**). Inactive RelA/p50 is held in the cytosol by its inhibitor IκBα until IκBα’s stimulus-signaled degradation frees the nuclear localization signal of RelA/p50, allowing it to translocate to the nucleus and seek out hundreds of target sites within our genome for activation (5, 6). By binding to target DNA sequences located within the *NFKBIA* gene promoter region, RelA/p50 also initiates production of its own inhibitor, IκBα, creating a negative feedback loop and facilitating the eventual stripping of RelA/p50 from DNA to cease activity (7–10). Due to its ability to influence the transcription of a variety of genes with roles in inflammation, immune responses and cell survival, the dysregulation of RelA/p50 is linked to inflammatory diseases, neurodegenerative diseases and cancer (3, 11).

**Figure 1.**
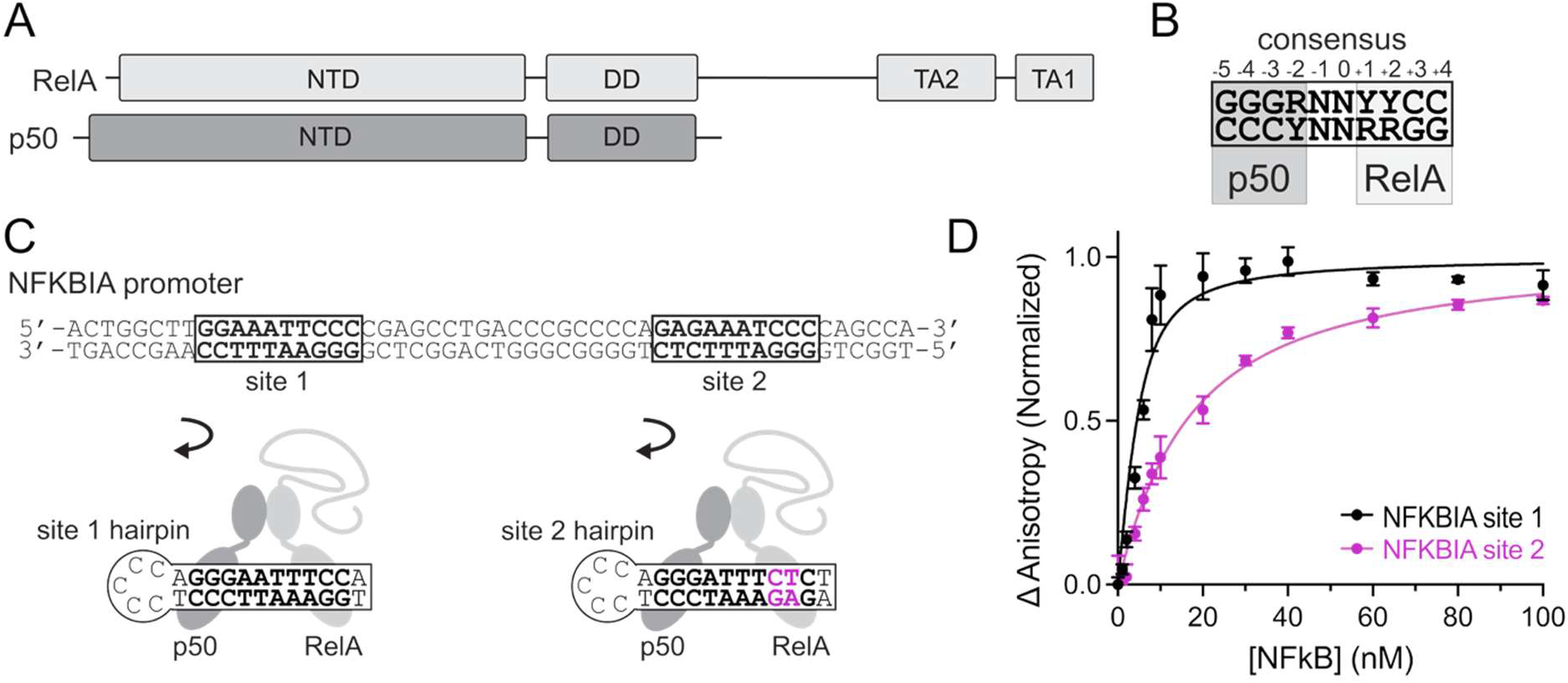
The NFκB heterodimer RelA/p50 binds to consensus κB and partial consensus (half-site) DNA sequences. (A) Both RelA (light gray) and p50 (dark gray) have an N-terminal domain (NTD) and a dimerization domain (DD), connected by a flexible linker. Additionally, RelA has a disordered transcription activation domain that includes two regions critical for protein-protein interactions – TA1 and TA2. (B) The RelA/p50 consensus κB site is a 10-base pair DNA sequence that can be separated into four-base pair RelA- and p50-binding regions with two base pairs in between. R denotes a purine, Y a pyrimidine, and N is any nucleotide. The binding sequence is numbered from -5 to +4 starting from the first p50-interacting nucleotide pair. (C) The NFKBIA promoter region contains multiple RelA/p50 binding sites, including site 1 (a full consensus kB site) and site 2 (a partial consensus kB site, or half-site). Site 2 nucleotides that are inconsistent with the consensus sequence are noted in magenta and occupy RelA-interacting positions +2 and +3. We investigated RelA/p50-DNA interactions using NFKBIA site 1 and site 2 sequences incorporated in DNA hairpins. (D) A representative binding experiment shows normalized change in anisotropy from fluorescein-labeled hairpin DNA plotted vs. RelA/p50 concentrations ranging from 0 to 100 nM. NFKBIA site 2 (magenta) has reduced binding affinity compared to site 1 (black). Error bars are standard deviation of change in anisotropy for each concentration prepared in triplicate. Mean K_D_ values and statistics from three such experiments are reported in **Table 1**.

**Table 1.**
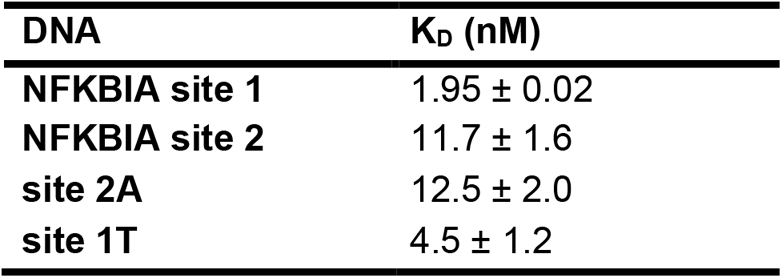
Binding affinities of RelA/p50 for NFKBIA sites 1 and 2 and hybrid DNA sequences. Mean K_D_ values from three independent experiments are reported with standard deviation.

| **DNA** | **K<sub>D</sub> (nM)** |
| --- | --- |
| **NFKBIA site 1** | 1.95 ± 0.02 |
| **NFKBIA site 2** | 11.7 ± 1.6 |
| **site 2A** | 12.5 ± 2.0 |
| **site 1T** | 4.5 ± 1.2 |

NFκB is a DNA binding protein, but it has an immunoglobulin-like fold instead of the more common structural categories of DNA binding proteins such as helix-turn-helix, helix-loop-helix, zinc-binding and leucine zipper (12). These common motifs are overwhelmingly α-helical, whereas NFκB uses the loops in its multi-domain structure to contact DNA (13). NFκB therefore may have the potential for a more flexible protein-DNA interaction. Indeed, our lab previously showed that whether free or bound to DNA, RelA/p50 is surprisingly dynamic (14, 15).

Such flexibility in the NFκB-DNA interactions naturally raises the issue of NFκB’s DNA sequence specificity. The RelA/p50 consensus sequence (referred to as a κB site) is 5’-GGGRNNYYCC-3’, where R denotes a purine, N denotes any nucleotide, and Y denotes a pyrimidine (**Figure 1B**) (16). The linkers and N-terminal domains (NTDs) of RelA/p50 interact at separate ends of this sequence – the p50 NTD with the 5’-GGGR sequence and RelA NTD with the YYCC-3’. Crystal structures of DNA-bound RelA/p50 have provided a detailed picture of extensive interactions occurring between the loops of NFκB and DNA in the complex. Binding is mediated by electrostatic interactions between basic residues of RelA/p50 and the negatively charged backbone of the DNA duplex as well as base-specific interactions, primarily within the major groove of DNA, that confer its DNA sequence specificity. The dimerization domains, linkers, and N-terminal regions of p50 and RelA create a binding pocket within which these DNA interactions occur (**Figure 1A and B**) (13, 17, 18). The result is high affinity binding with a dissociation constant in the low nanomolar range for the most ideal DNA sequences (19). Notably, however, although the backbone structure of RelA/p50 shows minimal changes when comparing the multiple crystal structures of consensus DNA-bound RelA/p50, amino acid side chains responsible for base-specific contacts have been shown to adopt different orientations across the different structures, and in some cases have been observed to engage in interactions with different nucleotide bases. For example, p50 residue R56 was observed to form bidentate contact with G^-4^ in one published structure but alternatively contacted G^-5^ in another. E60 was captured interacting with nucleotides at position -3 or -4, and the corresponding RelA residue E39 was observed to contact either the C^+3^ or the paired G^+3^ on the opposite strand (17).

In addition to genomic sites that match the full consensus sequence, there are also partial consensus sequences known as κB ‘half sites’ to which NFκB also binds. How RelA/p50 structural dynamics are impacted by binding DNA half-site sequences has not been studied, and it remains to be known whether the two subunits “read” DNA binding site halves independently or if both subunits have altered DNA interactions on partial consensus sequences. Because there are estimated to be many more half-sites in our genome (∼2 x 10^6^) compared to full κB sequences – (∼5 × 10^4^) it is critical to understand the nature of RelA/p50 interactions with half-sites (16). Ours and others’ previous measurements of DNA binding affinity found that half-sites bound 2- to 6-fold more weakly than the highest affinity consensus sequence (19–21). These typically contain the same 5’-GGGR motif – the p50-interacting end of the sequence – but do not match the consensus in the second half of the 10-base-pair sequence. For example, the *NFKBIA* promoter contains site 1, a full consensus κB sequence with the sequence 5’-GGGAATTTCC-3’, and site 2, a half site with the sequence 5’-GGGATTTCTC-3’ (**Figure 1C**) (22). Note that when site 2 is bound by the RelA/p50 heterodimer, the p50-interacting end of the sequence remains unchanged, whereas the RelA-interacting sequence strays from the consensus. In a crystal structure of RelA/p50 bound to the urokinase plasminogen activator (uPa) promoter, a half-site sequence that deviates from the consensus at +1 and +3 positions, the same p50 residues were observed to contact DNA vs. the consensus DNA-bound structures, but the exact interactions were subtly altered. For example, R56 contacted G^-4^ at one position instead of forming a bidentate interaction. In the RelA subunit, the nonconsensus +3 base pair resulted in R33 not contacting DNA (23).

To understand the RelA/p50-DNA interactions and protein structural dynamics when bound to κB half-sites, we used hydrogen deuterium exchange mass spectrometry (HDX-MS) to compare RelA/p50 bound to the two κB sites within the promoter region of the IκBα gene: *NFKBIA* site 1, a consensus κB site, and *NFKBIA* site 2, a half-site. We expected to observe a perturbed RelA DNA-interaction but an unchanged p50-DNA interaction, but our results instead show a coordinated conformational rearrangement of both subunits’ NTDs and linkers when bound to half-site DNA. We explored possible mechanisms of this coordination using all-atom molecular dynamics (MD) simulations and observed that DNA adopts a unique position within the RelA/p50 pocket when bound to a half-site vs. full consensus sequence, propagating a change in structural dynamics between the NTDs. Our results help to explain how RelA/p50 can adaptably bind to many different DNA sequences with high affinity and activate hundreds of genes.

## Results

### NFκB RelA/p50 binds with 6-fold higher affinity to *NFKBIA* site 1, a consensus site, compared to *NFKBIA* site 2, a half-consensus site

To explore how nuclear-localized NFκB might interact with the multitude of available consensus and half sites in our genome, we expressed and purified RelA_19-549_/p50_39-350_ (hereafter referred to as RelA/p50) and carried out fluorescence anisotropy experiments using hairpin DNA containing either *NFKBIA* site 1 – a consensus NFκB DNA sequence – or *NFKBIA* site 2 – a half-consensus sequence (**Figure 1C**). Change in fluorescence anisotropy over a range of RelA/p50 concentrations was used to determine the binding dissociation constant (K_D_) for *NFKBIA* site 1 and site 2, with representative binding curves for each shown in **Figure 1D**. Summary K_D_ results and statistics, reported in **Table 1**, report that the measured K_D_ of RelA/p50 for site 1 was 1.95 ± 0.02 nM whereas for site 2 was 11.7 ± 1.6 nM. We hypothesized that this change in affinity between site 1 and site 2 binding is due to a reduction in the number of sequence-specific interactions that form between RelA and its interacting DNA sequence because of the swapped +2 and +3 nucleotides. We expected the p50 interactions to be the same between site 1 and site 2 DNA because both half-sequences match the consensus (**Figure 1B**).

### HDX-MS probes structural dynamics of RelA/p50 bound to consensus and half-site DNA

To gain insight into the dynamics of the flexible loops of NFκB that engage DNA corresponding to a consensus vs. half-site sequence, we carried out HDX-MS with RelA/p50. We compared free RelA/p50 to RelA/p50 bound to either *NFKBIA* site 1 or site 2 in the form of hairpin DNA containing a single 10-base pair target site. Deuterium exchange was carried out for 0, 0.5, 1, 2 and 5 minutes, a range of incubation times that report on protein dynamics in the biochemically relevant microsecond to millisecond time regime (24), and backbone amide deuterium uptake was measured by mass spectrometry. The deuterium exchange of regions of RelA/p50 outside of the DNA-binding pocket remained the same across all conditions indicating that DNA binding, regardless of sequence, did not impact the overall protein structure (**Supporting Figure 1**). However, we identified significant sequence-dependent differences in exchange in regions within the DNA-binding pocket that shed light on how RelA/p50 accommodates half-site vs. full consensus sequences.

### Flexible linkers and a subset of N-terminal domain loops of RelA and p50 are highly sensitive to consensus vs. half-site binding

In previously-resolved crystal structures, the flexible linkers that connect the NTDs and dimerization domains of RelA/p50 are observed to play a direct role in DNA binding, with side chains inserted into the major groove of DNA interacting with specific bases: in RelA, arginine 187 interacts with T^+2^ and in p50, the less-restrained lysine 241 has been shown to contact nucleotide bases in the 0, -1 and -2 position (17). After 30 sec. of labeling time in our HDX-MS experiments, the flexible linker of RelA had 3.5 Da less deuterium uptake when site 1-bound compared to free protein, indicating that the presence of bound consensus DNA significantly protects this linker from exchange. The same residues were much less protected upon site 2 binding (only 1 Da decrease vs. free protein) (**Figure 2A**). This discrepancy between site 1 and 2-bound protein is evidence of the DNA sequence sensitivity of the RelA linker. This difference was expected due to the non-consensus RelA-interacting residues (T^+2^ becomes C^+2^) in the half-site DNA sequence.

**Figure 2.**
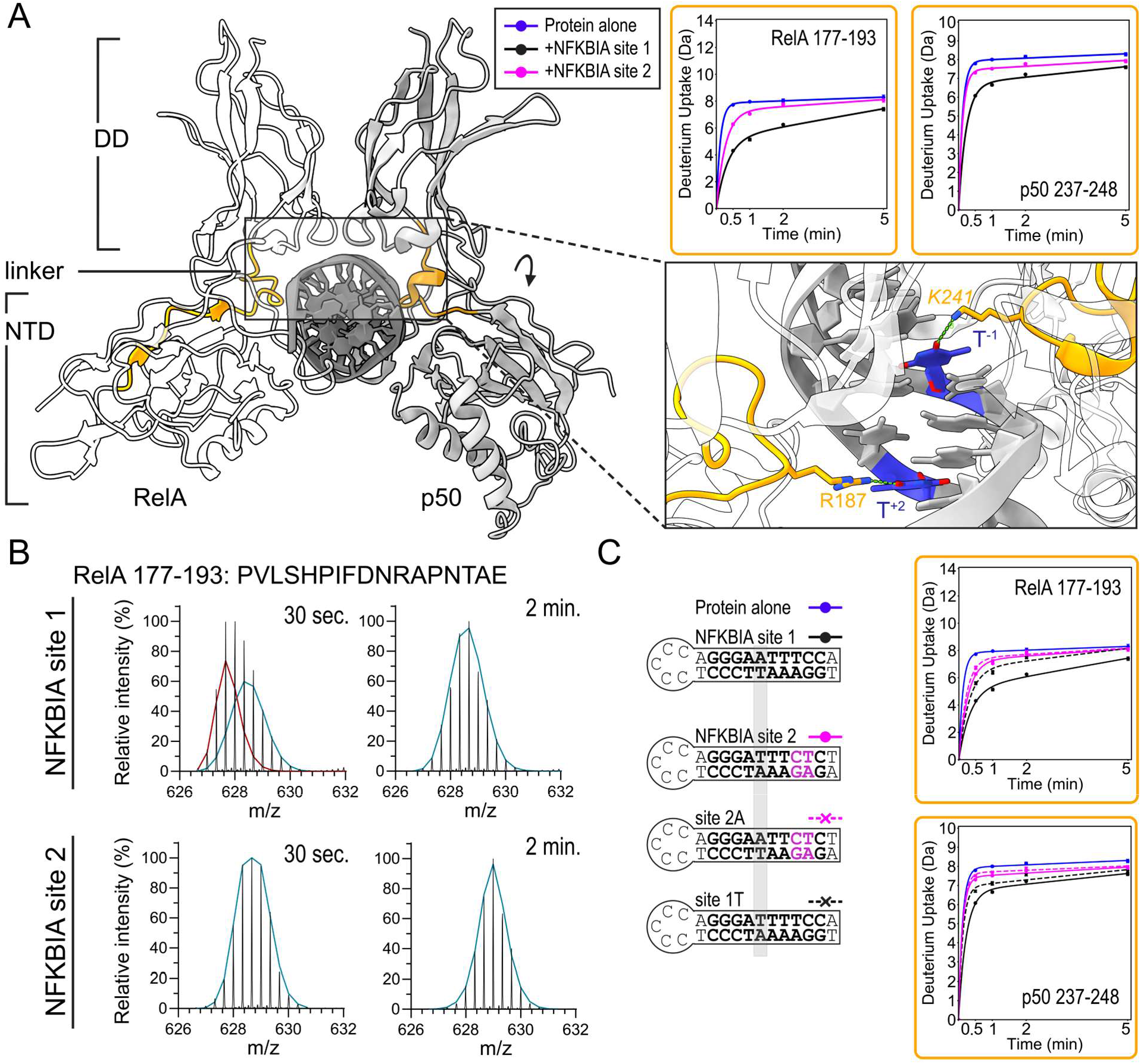
HDX-MS reveals DNA sequence-dependent changes in structural dynamics for both flexible linkers of RelA/p50. Hydrogen-deuterium exchange mass spectrometry (HDX-MS) was used to analyze deuterium uptake of RelA/p50 alone vs. bound to hairpin DNA containing *NFKBIA* site 1 and hairpin DNA containing *NFKBIA* site 2. (A) Deuterium uptake from HDX-MS analysis of free (blue) vs. site 1-(black) and site 2-bound (magenta) RelA/p50 was quantified for regions that included flexible linkers of RelA (left, white) and p50 (right, gray) and mapped onto PDB 1LE5 (17) in yellow. Subunit identity and residue range are included on the right side of each plot. For both linker regions, site 1 binding resulted in significantly lower deuterium uptake compared to site 2 binding. Zoomed region (90° rotated top-down view) of PDB 1LE5 identifies RelA linker residue R187 and p50 linker residue *K241* as forming specific interactions with T^+2^ and T^-1^ of a consensus κB site. NFKBIA site 1 has a T/A pair at position +2, while NFKBIA site 2 instead has C/G pair at the +2 position (Figure 1C). (B) Relative intensity vs. m/z is shown for site 1-bound and site 2-bound RelA region 177-193 (sequence displayed with R187 in bold) after 30 sec. and 2 min. of deuterium labeling. RelA region 177-193 was analyzed for bimodal character using HX Express v3. After 30 sec. of labeling, region 177-193 did not fit a single population mass envelope when bound to site 1; the m/z distribution was best fit with two distinct populations (lower m/z population in red, higher m/z population in blue), whereas site 2 was best fit with one population (blue). By 2 minutes of labeling, site 1- and site 2-bound RelA linker resolved into a single deuterated population (2 min., blue). (C) Impact of changing the middle (-1 position) nucleotide pair was explored using hybrid DNA hairpins with the -1 position highlighted in gray. The sequence of site 2A (magenta, dashed line) matched NFKBIA site 1 at position -1 and otherwise matched site 2, while the sequence of site 1T (black, dashed line) matched NFKBIA site 2 at position -1 and otherwise was identical to NFKBIA site 1. Site 2A-bound RelA/p50 had identical deuterium uptake vs. the NFKBIA site 2-bound form through both linker regions, while site 1T binding resulted in less protection through both linkers as compared to site 1 binding.

For the site 1-bound RelA linker region, the deuterated mass envelope could be resolved into two distinct populations at the 30 sec. timepoint that merged into a single detectable population by the 2 min. timepoint (**Figure 2B**). This bimodal character suggests distinct states that undergo slow interconversion on the timescale of deuterium exchange (25). Importantly, only RelA bound to site 1 showed the more protected population, whereas when bound to site 2, only the higher-exchanging population was observed. This led us to hypothesize that the more protected (less exchanging) of these populations corresponds to a hydrogen-bonded R187 residue as has been observed in crystal structures (**Figure 2A**) (17), an interaction that would not be present upon site 2 binding because of the swapped sequence at the +2 position.

Surprisingly, the p50 linker also showed altered deuterium uptake upon binding site 1 vs. site 2 DNA. This linker experienced a 2 Da decrease in deuterium uptake after 30 sec. of deuterium labeling when bound to site 1 as compared to free protein, and only 0.5 Da less uptake when bound to site 2, also indicating DNA sequence sensitivity (**Figure 2A**). This was contrary to our expectation, since p50 interacts with a consensus half-sequence when binding either site 1 or site 2. To determine if it could be explained by a change in the middle base pair (-1 position) between site 1 and site 2, despite this being a supposed nonspecific nucleotide, we created a hybrid sequence (called site 2A), where the -1-nucleotide position was mutated to match site 1. Site 2A isolates sequence changes to specific bases with which RelA interacts and maintains the same sequence bound by p50 as site 1. RelA/p50 bound this hybrid half-site sequence with the same affinity as NFKBIA site 2 (**Table 1**). Comparing the deuterium uptake resulting from site 2 binding vs. site 2A binding revealed no difference (**Figure 2C**). The observation that p50 showed different protection when bound to site 1 vs. site 2A, which have identical p50-interacting sequences (5’-GGGAA), indicates that changes in deuterium uptake by p50 upon binding half-site DNA are in response to the altered RelA-DNA interaction rather than changes in the DNA bases p50 directly contacts.

We created a second hybrid site that changes the -1 position in the site 1 sequence to match that of site 2, which we called site 1T. This sequence remains a full consensus κB sequence and has an identical RelA-interacting sequence to site 1. Notably, this sequence caused increased deuterium uptake in the linker regions of both RelA and p50 compared to wild-type site 1 (**Figure 2C**). This suggests that RelA linker dynamics can also respond to altered dynamics in the p50 linker, as the RelA-interacting sequence is identical between these two sequences, and the RelA linker has not been shown to directly interact with position -1 in the binding sequence. The altered linker dynamics in p50 could be explained by the change in the -1 nucleotide. This change in linker dynamics also corresponded to a slightly higher K_D_ for site 1T vs. site 1, which at 4.5 nM fell in between the binding affinity of site 1 and sites 2/2A (**Table 1**).

We analyzed the NTD loops of RelA/p50 that contact the DNA-binding cavity and found that some showed differences in protection depending on which DNA sequence was bound and others did not (**Figure 3**). RelA residues 120-132 exhibited less protection with site 2/2A binding vs. site 1/1T, whereas RelA loop residues 23-34 experienced equal deuterium uptake regardless of which DNA sequence was bound. In the p50 subunit, residues 43-55 were significantly more protected when bound to site 1 or 1T DNA compared to when bound to site 2/2A. Residues 56-86 of p50 also exhibited DNA sequence-dependent protection, while residues 135-151 had equivalent decreases in deuterium uptake with each site. The observation that linkers and some loops of both subunits experience DNA-sequence dependent HDX-MS changes, despite a p50-interacting half sequence that matched the consensus, provides evidence for the coordinated adjustment of both subunits on a DNA half-site as opposed to each subunit independently “reading” and conforming to a separate half-consensus sequence. We did not observe any differences in deuterium uptake in NTD loop peptides upon binding site 1 vs. 1T or site 2 vs. 2A DNA. This suggests that changes in structure and dynamics due to the nucleotide identity at the -1 position are limited to the linker peptide and do not extend into the NTDs.

**Figure 3.**
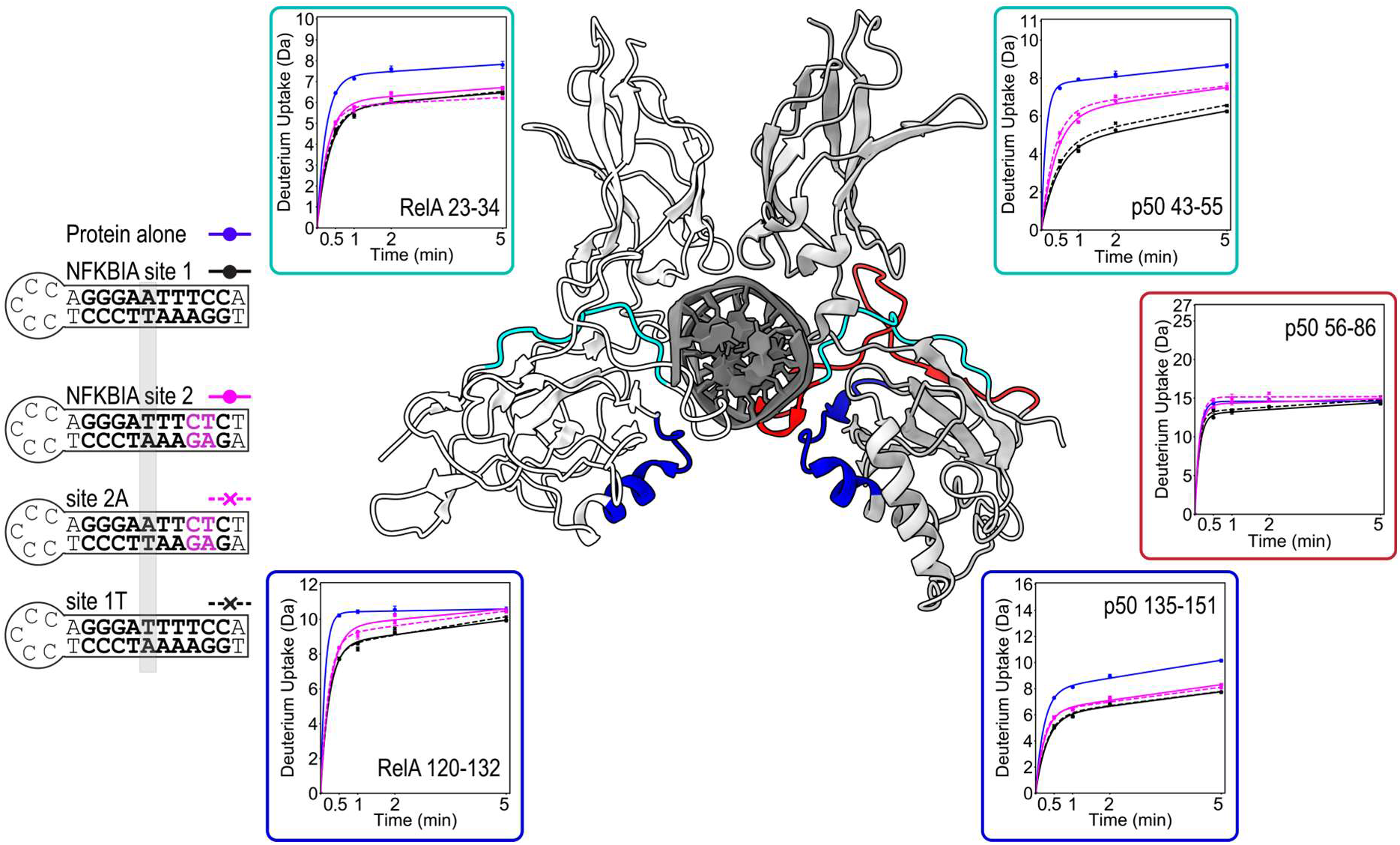
Unexpected dynamics at the p50 NTD are not explained by the middle (-1) nucleotide identity of site 2. Deuterium uptake from HDX-MS analysis of free (blue) vs. site 1-(black) and site 2-bound (magenta) RelA/p50 was compared to hybrid sites 2A (magenta dashed line) and 1T (black dashed line) for regions in the NTDs of RelA (left) and p50 (right). Hybrid site hairpin sequences are compared on the left, with the -1-position highlighted in grey. Nucleotides that deviate from the consensus sequence in sites 2 and 2A are noted in magenta. Throughout the RelA/p50 NTDs, hybrid site 1T binding resulted in uptake that was identical to site 1 binding, and site 2A-bound RelA/p50 uptake was identical to site 2-bound RelA/p50. Uptake plots are color-coordinated with protein regions mapped onto PDB 1LE5.

### Dimerization domains of RelA/p50 are involved in DNA binding but generally insensitive to DNA sequence

We initially hypothesized that the sequence sensitivity of p50 was the result of RelA inducing changes in the structural dynamics of p50 via a pathway through the dimerization domains (DDs), which form the only point of direct contact between the RelA and p50 subunits. To explore this possibility, we examined the impact of consensus vs. half site binding on DNA-interacting DD loops and the DD interface. The top of the DNA binding pocket, formed by multiple DD loops of RelA/p50, showed significant protection upon DNA binding (**Figure 4A**). Regions of RelA that include basic residues K218, K221, and R246, as well as regions with corresponding residues in p50 (*K272, K275 and R305*) experienced 0.6-1.5 Da protection when DNA-bound. The deuterium uptake for these RelA DD regions was the same regardless of whether site 1 or site 2 DNA was bound. The results were similar for the p50 DD loops, however there was a small divergence in site 1 vs. site 2 deuterium uptake for region 272-279 at the later timepoints. This again suggests that changes in the RelA-DNA interactions are propagated to p50, although the result is most prominent in the NTD. Overall, these data show that the DD loops forming the top of the DNA binding pocket are generally insensitive to the differences in DNA sequence. Similar results were obtained for the DD interface. This region showed no uptake change upon DNA binding (**Figure 4B**). The observation that the DDs and their interface are insensitive to DNA sequence variation indicates that the changes in the linkers and NTDs when RelA binds a different DNA half-sequence are likely not propagated to p50 through the directly interacting DDs.

**Figure 4.**
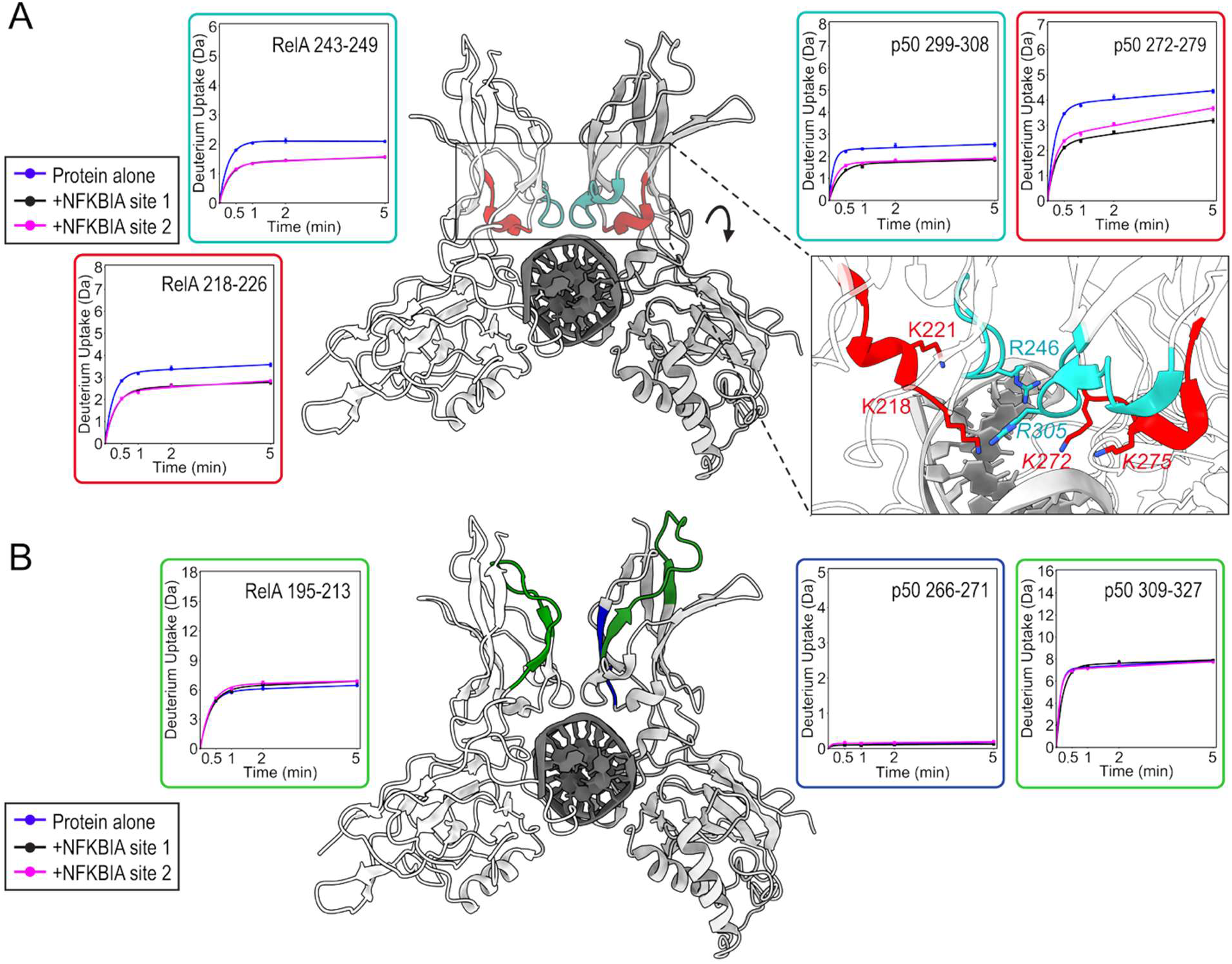
Dimerization domain structural dynamics are not affected by DNA sequence. Deuterium uptake from HDX-MS analysis of free (blue) vs. site 1-(black) and site 2-bound (pink) RelA/p50 was quantified for (A) DNA-interacting dimerization domain loops of RelA (left) and p50 (right) and (B) dimerization domain interface regions of RelA and p50. In both cases, regions of interest are mapped onto PDB 1LE5 and color-coordinated to uptake plot outlines. The deuterium uptake of dimerization domain regions generally did not show significant changes depending on DNA sequence. The inset in (A) shows basic residues that are oriented toward DNA in dimerization domain loops of RelA and p50 (p50 residues in italics).

### Molecular dynamics simulations analyzed by HDXer capture the sequence sensitivity of NFκB-DNA binding

To investigate the conformational dynamics of the protein-DNA complex, we carried out three 7-8 µs all-atom MD simulations comparing RelA/p50 in complex with site 1 vs. the hybrid site 2A hairpin DNA. Comparing these two sites allowed us to investigate the impact of sequence changes isolated to RelA-interacting nucleotides on both RelA and p50. We first compared the root mean square fluctuation (RMSF) values for the site 1 and site 2A DNA-bound structures to evaluate whether flexibility is responsible for the differences observed in the HDX-MS results. The simulations identified multiple regions where site 2A-bound RelA/p50 residues had mostly higher RMSF values compared to those of site 1-bound RelA/p50, with changes concentrated in the NTDs and linkers of both subunits (residues 19-194 of RelA, and 39-248 of p50). In agreement with the HDX-MS data, the dimerization domains (residues 249-350 of p50 and 195-291 of RelA) showed minimal change in average RMSF when assembled on site 2A vs. site 1 (**Supporting figure 2**).

To make direct comparisons between MD simulations and experimental HDX-MS, we used HDXer to generate predicted deuterium uptake from the simulations for each RelA/p50 peptide identified in HDX-MS experiments (26). The HDXer-predicted deuterated fractions from simulations matched well with the experimental results for RelA bound to site 1 and site 2A (**Supporting figure 3**) and for p50 bound to site 1 and site 2A (**Supporting figure 4**). For RelA, the difference in predicted deuterium uptake between site 1 and site 2A was consistent with experimental results – specifically the linker region (residue 185-194) was predicted to exchange more with site 2A binding than with site 1 (**Supporting figure 5A**). For the p50 linker region (residues 237-248), however, the HDXer-predicted difference between site 1 and site 2A from simulations was opposite of experimental results, predicting that the linker would exchange more when bound to site 1 vs. site 2A (**Supporting figure 5B**). To construct an ensemble of conformations from our starting simulations that more accurately matched experimental HDX-MS data, we used HDXer to carry out HDX ensemble reweighting (**Supporting figures 3 and 4**) (26). While the predictions of deuteration from reweighted simulations did not fully recapitulate the trend observed in the p50 linker by HDX-MS experiments, where site 2A binding resulted in greater deuterium exchange vs. site 1 binding, it did minimize the difference compared to the prediction from unweighted simulations (**Supporting figure 5**).

The ΔRMSF values for the reweighted simulations were again calculated to evaluate changes in flexibility across RelA/p50 depending on DNA sequence (**Figure 5A**). Both the unweighted and reweighted simulations captured an increase in RMSF in p50 NTD residues 150-230 with site 2A binding vs. site 1, as well as residues in the RelA linker and NTD, consistent with the decrease in binding affinity of site 2A DNA (**Table 1**) (**Figure 5A, Supporting figure 2**). The simulations therefore observed an overall higher flexibility within both RelA and p50 when bound to the half-site DNA as compared to the full consensus DNA, despite a consistent p50-interacting sequence (5’-GGGAA), in agreement with experimental HDX-MS results.

**Figure 5.**
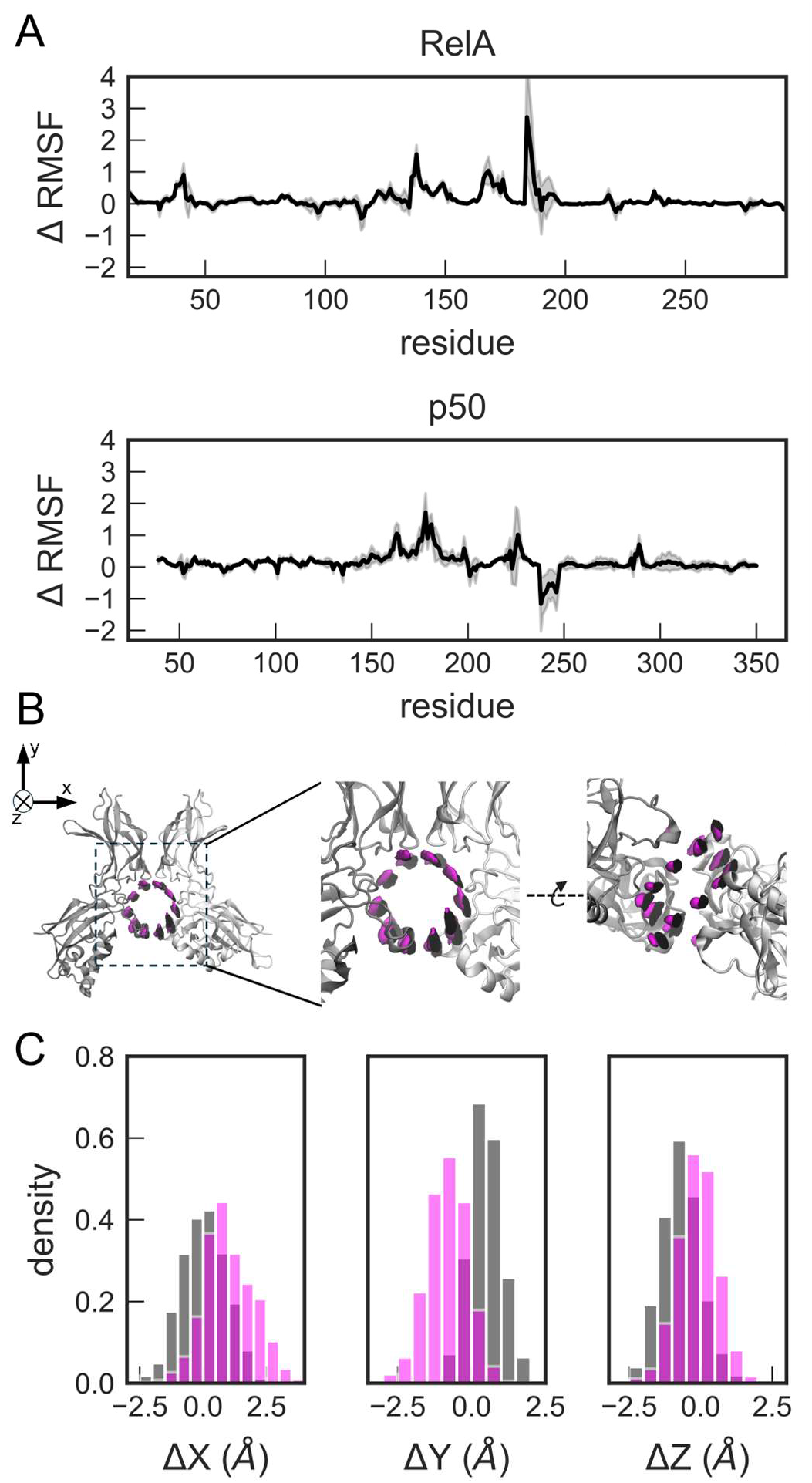
Half-site DNA adopts a unique position within the RelA/p50 binding pocket. (A) Changes in the RMSF of RelA and p50 residues when bound to NFKBIA site 2A vs. site 1 (RMSF_site 2A_ − RMSF_site 1_ = ΔRMSF) were calculated from HDXer reweighted simulations (26). The standard error is shown as a gray envelope. (B) Density of the three-dimensional distributions of the DNA phosphorus atoms contoured at 0.2% occupancy per Å^3^ show the unique positions occupied by site 1 vs. 2A DNA. (C) To quantify this difference, net displacement in x, y and z directions of each DNA molecule relative to DNA binding loops of RelA/p50 are displayed in density plots. For panels B and C, site 1 is in black and site 2A is in magenta.

### Molecular dynamics simulations capture a shift in the position of half-site DNA within the binding pocket of RelA/p50

To analyze the global movement of DNA relative to RelA/p50, we analyzed each set of trajectories (site 1 DNA-bound vs site 2A DNA bound) by first aligning the phosphorus atoms of the base-pairing residues of the initial structures to an idealized B-DNA double helix whose superhelical axis was aligned to the z-axis. Then, the rest of the trajectory frames were aligned to these initial, z-aligned structures by aligning the loops that contact the DNA (see Methods for more details) (**Figure 5B**). Using this strategy, we uncovered a measurable shift in the average displacement of the RelA/p50 protein relative to site 1 DNA vs. site 2A DNA throughout the simulations, most significantly along the x- and y-axes, resulting in distinct positions of the two DNA molecules within the binding pocket of RelA/p50 (**Figure 5C**). The average positions of the phosphorus atoms shifted by 0.97 Å in x, -1.26 Å in y and 0.59 Å in z in the site 2A simulations as compared to the site 1 simulations. Overall, the DNA shifted closer to p50 in the site2A simulations as compared to site 1. Because DNA makes sequence-specific contacts with both NTDs and linkers of RelA/p50, even a modest shift in its position would impact interactions with both subunits. Taken together, HDX-MS, RMSF differences and DNA position analysis demonstrate the impact of half-site vs. consensus DNA binding on the conformational flexibility of the overall protein-DNA complex.

## Discussion

RelA/p50 has been observed *in vivo* to bind DNA lacking a full consensus κB site (27–29), including DNA that lacks even a half-consensus sequence (28). Based on crystal structures, RelA residues form ∼three base-specific hydrogen bond(s) with the +2 and +3 nucleotide pairs (R33-G^+3^, E39-C^+3^, R187-T^+2^, PDB 1LE5) (17). Free energy calculations based on the approximation that each H-bond contributes 1 kcal/mol of stability to the complex (30) show that the loss of three H-bond interactions when the half-site sequence is bound would be expected to decrease binding affinity by ∼160 fold. However, consistent with prior results, we found that the half-consensus site DNA binding affinity was only 6-fold weaker compared to the full consensus (19). To understand how NFκB could retain high affinity binding to half-site DNA, we carried out HDX-MS and all-atom MD simulations. Our results shed light on how the dynamic structure of RelA/p50 bound to DNA can adjust to bind with high affinity to both full consensus and half-consensus DNA sequences.

We found that when the RelA-binding sequence no longer corresponds to the κB consensus, changes in amide exchange were observed in the linkers and the NTD loops of both RelA and p50, suggesting a coordinated adjustment of RelA and p50 to achieve half-site DNA binding, as opposed to independent DNA interactions with each subunit. These results indicate that the p50 subunit adjusts its conformational ensemble even when it is bound to the same DNA half-sequence. We initially hypothesized that the p50 conformational adjustment would occur because the RelA conformational change upon binding a non-consensus sequence would be transmitted through the dimerization domains to p50. However, the lack of DNA sequence effect on dimerization domain dynamics indicated that communication between the RelA and p50 subunits did not involve the dimerization domains. Instead, all-atom MD simulations showed a change in the position of DNA within the RelA/p50 DNA binding pocket. Because DNA connects both subunits and forms sequence-specific contacts with RelA and p50, a shift in the position of the DNA would alter the protein-DNA interactions in both subunits. The structural rearrangements of RelA and p50 are therefore observed in the linkers and NTDs which make specific contacts with DNA bases, but not in the dimerization domains which make primarily nonspecific interactions with the DNA backbone.

The ability of RelA/p50 to adapt its structure to different DNA sequences can be largely attributed to the lack of secondary structure in its DNA-binding regions; NFκB binds DNA via loops instead of the more common α-helices present in other DNA binding proteins (12). As we noted earlier, previous structural analysis of RelA/p50 identified that many residues were able to interact with more than one nucleotide (17). More recently, Li et al. made the interesting observation that, in the RelA homodimer, the equilibrium between side chain conformations was modulated by the identity of the middle, supposedly nonspecific nucleotide pairs in the κB sequence (31). We also observed the importance of the middle nucleotides. A conservative single A/T to T/A swap of the -1 nucleotide pair (site 1 vs. site 1T), which is nonspecific in the defined consensus sequence, caused altered dynamics through the flexible linkers of both RelA and p50 and a measurable decrease in binding affinity. Our results provide strong evidence that NFκB can bind with high affinity to many different DNA sequences by adjusting the conformational ensemble of DNA-interacting loops and a coordinated optimization of the DNA position between the two subunits of the dimer. These results help explain how NFκB maintains high affinity for many different DNA sequences, even when they differ in sequence at positions contacted specifically by NFκB.

## Methods

### Protein expression and purification

N-terminally tagged murine 6xHis-p50_39-350_ and untagged murine RelA_19-549_ were co-expressed from a pET11a vector containing both genes as described previously (32). The vector was transformed into *E. coli* BL21(DE3) cells and grown at 37°C with ampicillin selection until an OD_600_ of 0.6-0.8 in M9-ZN media. After cooling cultures on ice, IPTG was added at 0.2 mM to initiate protein expression, and cultures were incubated in a shaker for 16 hours at 18°C. Following expression, cells were harvested by centrifugation and resulting pellets were stored at -80°C.

Purification of 6xHis-p50_39-350_RelA_19-549_, hereafter referred to as RelA/p50, was carried out as previously described (19). All purification steps took place at 4°C. In brief, cells were lysed using sonication and lysate was clarified by centrifugation before being applied to a Ni^2+-^NTA gravity column. The elution was dialyzed overnight to remove imidazole, separated into aliquots and stored at -80°C. Immediately before experiments were carried out, protein aliquots were thawed and purified using cation exchange chromatography (MonoS, GE healthcare) with a 0-700 mM NaCl gradient. Eluted fractions containing RelA/p50 were further purified using size exclusion chromatography (Superdex 200 column, GE healthcare) in NFκB SEC buffer (25 mM Tris, 150 mM NaCl, 0.5 mM EDTA, 1 mM DTT, adjusted to pH 7.5 at room temperature [RT]) to carefully separate full-length RelA/p50 from truncated protein products retained through previous purification steps. RelA/p50 protein purity was assessed using SDS-PAGE, and concentration was determined using 280 nm absorbance on a NanoDrop spectrophotometer. Protein was stored at 4°C before use.

### Fluorescence Anisotropy

The following 5’ FAM(fluorescein)-labeled hairpin DNA sequences were purchased from Integrated DNA Technologies:

NFKBIA site 1: 5’-T<u>GGAAATTCCC</u>TCCCCCA<u>GGGAATTTCC</u>A-3’
NFKBIA site 2: 5’-A<u>GAGAAATCCC</u>TCCCCCA<u>GGGATTTCTC</u>T-3’
NFKBIA site 2A: 5’-A<u>GAGAATTCCC</u>TCCCCCA<u>GGGAATTCTC</u>T-3’
NFKBIA site 1T: 5’-T<u>GGAAAATCCC</u>TCCCCCA<u>GGGATTTTCC</u>A-3’

Labeled hairpin DNA was prepared for binding experiments by diluting to 100 nM in NFκB SEC buffer, heating to 95°C for 5 minutes and cooling slowly over 2-3 hours. Hairpin DNA was diluted to its final experimental concentration (5 nM) in NFκB SEC buffer with RelA/p50 at protein concentrations ranging from 0-100 nM, with each concentration prepared in triplicate in a black 96-well plate. Plates were incubated at RT for a minimum of 2 hours. Fluorescence anisotropy was measured at RT on a Molecular Devices SpectraMax iD5 Microplate Reader. The excitation and emission wavelengths were 485 nm and 535 nm, respectively. The integration time of data collection was 100 ms. The resulting anisotropy calculated using the equation r = [I(V,V) – GI(V,H)]/[I(V,V) – 2GI(V,H)] where r is anisotropy, I(V,V) is fluorescence intensity in the parallel direction, I(V,H) is fluorescence intensity in the perpendicular direction, and G is the grating factor. The G-factor used for calculating anisotropy was 1, as previously determined for this instrument.

Fluorescence anisotropy data for the hairpin DNA sequences were fit to the following equation to determine equilibrium binding affinity:

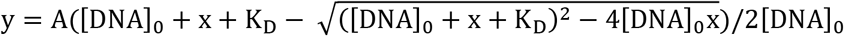

where y is the change in fluorescence anisotropy, A is the maximum change in fluorescence anisotropy, x is the varying concentration of RelA/p50, [DNA]_0_ is the concentration of DNA (5 nM), and K_D_ is the equilibrium dissociation constant (33). Three separate fluorescence anisotropy experiments were carried out on three separate days from distinct preparations of RelA/p50 and hairpin DNA. For each assay, data were fit to the aforementioned equation to determine the K_D_. Due to slight variability in the plateau anisotropy value between experiments, data from each experiment was normalized by the value of A determined for that experiment in order to generate plots like the representative example shown in **Figure 1D**. Mean and standard deviation of the measured K_D_ values were calculated from the three replicates for each condition.

### HDX-MS

The four hairpin DNA sequences described above were obtained from Integrated DNA Technologies without the 5’ (FAM)fluorescein label for use in HDX-MS analysis of RelA/p50-DNA interactions. For HDX-MS, hairpin DNA was prepared by diluting to 100 µM in NFκB SEC buffer, heating to 95°C for 5 minutes and cooling slowly. Hairpin DNA (10 µM final) was mixed with RelA/p50 (5 µM final) in NFκB SEC buffer; a 1:2 heterodimer:hairpin DNA ratio was used to achieve fully DNA-bound RelA/p50 based on binding affinities measured by fluorescence anisotropy. DNA-free samples of RelA/p50 were also prepared.

HDX-MS experiments were carried out using a Waters Synapt G2Si time-of-flight mass spectrometer with a nanoACQUITY UPLC system with HDX technology and a LEAP autosampler as in (19). Prepared samples were stored in a 1°C chamber for the duration of the experiment. For each timepoint, 4 µL of sample was equilibrated to RT, mixed with 56 µL of D_2_O buffer (25 mM Tris, 150 mM NaCl, 1 mM DTT, and 0.5 mM EDTA, pH 7.5 at RT), and incubated for the desired time. Timepoints were collected in triplicate for 30, 60, 120 and 300s. Triplicates of a 0s timepoint were collected using H_2_O buffer that was otherwise identical in composition. Following incubation, 50 µL of sample solution was mixed with 60 µL of quench solution (3 M Guanidine-HCl, prepared with formic acid so that final sample-quench mixture achieved pH 2.8 on ice). The quenched sample was then injected onto an in-line pepsin column (immobilized pepsin, ThermoFisher, Inc). The resulting peptides were trapped and separated on a C18 column (Acquity UPLC BEH C18, 1.7 µM, 1.0 x 50 mm; Waters Corporation) using a 7% to 85% acetonitrile gradient with 0.1% formic acid over 7.5 min and directly electrosprayed into the mass spectrometer. The mass spectrometer was set to collect data in the Mobility ESI+ mode, with mass acquisition range of 200 to 2000 (m/z) and scan time of 0.4 s. Continuous lock mass correction was accomplished with an infusion of leuenkephalin (m/z = 556.277) every 30s (mass accuracy of 1 ppm for calibration standard). For peptide identification, the mass spectrometer was set to collect data in MSE, mobility ESI+ mode instead. The peptides were identified from triplicate of MSE analyses of 10 µM protein solution, and data were analyzed using PLGS 2.5 (Waters Corporation). The peptides identified in PLGS were then analyzed in DynamX 3.0 (Waters Corporation).

The relative deuterium uptake for each peptide was calculated by comparing the centroids of the mass envelopes of the deuterated sample vs. the undeuterated controls following previously published methods (34). The deuterium uptake was corrected for back exchange as previously described (35). Deuterium uptake plots were generated in DECA (github.com/komiveslab/DECA), and the data are fitted with an exponential curve for ease of viewing (35). Deuterium uptake is reported in uptake plots with y-axes that range from 0 to the maximum theoretical uptake for each peptide. Peptides were analyzed for multimodal distribution using HX-Express v3 (25). Community guidelines have been followed, and the data are publicly available on the Massive data repository (36).

### MD simulations

Starting models were generated using Alphafold3 (37) to predict the atomic coordinates of the DNA-binding domains of NFκB RelA_19-291_/p50_39-350_ bound to the DNA hairpin sequences for site 1 and site 2A. Each model was solvated in an octahedral periodic box of optimal point charge water and 150 mM NaCl using AmberTools24 (38). The site 1 simulations are composed of 160,439 total atoms and 49,997 waters. The site 2A simulations are composed of 161,438 total atoms and 50,330 waters. Simulations were run at constant pressure (1 bar) and temperature (298 K) using the Monte Carlo barostat, the Langevin thermostat, a nonbonded cutoff of 10 Å, the ff19SB protein forcefield (39) and the OL24 DNA forcefield (40). Protonation states of histidines were assigned using H++ (41). Using the CUDA enabled version of Amber24’s pmemd (38), three parallel simulations for each model were equilibrated using the AmberMDprep protocol (42). Once equilibrated, the production simulations were run at 4 fs timestep with hydrogen mass repartitioning. For site 1, three simulations of 7 ms were run for an aggregate total of 21 ms. For site 2A, three simulations of 8 ms each were run, totaling 24 ms aggregate time.

To compare the HDX-MS data to the simulations, simulations were stripped of solvent and subsampled to one frame per 10 ns using cpptraj in AmberTools24. HDXer was used to compare HDX-MS data to the MD simulations (26). Within HDXer, reweighting of MD trajectories was performed with parameter optimization using a lambda value of 4.0 for site 1 and 5.0 for site 2A, which correspond to work values of 7.21360 kJ/mol and 7.42038 kJ/mol, respectively. To calculate the reweighted RMSF, each domain was decomposed for alignments and subsequent variance calculations. The six domains were split into the nucleotide binding domain for p50 (residues 39-237), the linker domain for p50 (residues 238-247), the dimerization domain of p50 (residues 248-350), nucleotide binding domain for RelA (residues 18-183), the linker domain for RelA (residues 184-194), the dimerization domain of RelA (residues 195-291). Each domain was aligned to the initial reference model, according to C_a_ RMSD, then the weighted RMSF was calculated as:

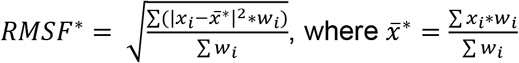

DNA phosphorus distributions were calculated first by aligning the phosphorus atoms of the base pairing residues of the initial structures were aligned to an idealized B-DNA double helix whose superhelical axis was aligned to the z-axis. This idealized B-DNA was generated using PyMol [8]. The rest the trajectory frames were aligned to these initial, z-aligned structures calculated by aligning based on C_a_ RMSD of protein residues contacting the DNA (residues 54, 56, 57, 59, 60, 62, 63, 64, 65, 74, 77, 113, 114, 136, 141, 143, 144, 145 for p50 and residues 28, 31, 32, 33, 35, 36, 39, 40, 41, 47, 51, 52, 53, 56, 122, 123, 124 for RelA). Once aligned, three-dimensional distributions were calculated with a bin spacing of 1 Å. The x, y, z shift distributions were also calculated from these aligned trajectories. For these calculations, a trajectory spacing of one frame per nanosecond was used.

## Data availability

The raw HDX-MS data files and analyzed data are available at massive.ucsd.edu username MSV000103724_reviewer, password a. MD simulation data is available at this zenodo link: 10.5281/zenodo.23677051.

## Acknowledgements

NEB acknowledges support from NIH grant 1R35GM165612. EAK acknowledges support from NIH grant R01 GM141182. TRG and LS acknowledge support from NIH T32 GM139795.

