## Supporting Data for "Optimization of the NFκB-DNA ensemble leads to high affinity binding of partial consensus sites"

Table S1. HDX-MS parameters

| **Data Set** | ***RelA/p50*** | ***RelA/p50* + site 1 DNA** | ***RelA/p50* + site 2 DNA** | ***RelA/p50* + site 1T DNA** | ***RelA/p50* + site 2A DNA** |
|---|---|---|---|---|---|
| HDX reaction details | 25 mM Tris-HCl, 150 mM NaCl, 0.5 mM EDTA 1 mM DTT, pH 7.5 @ 25 °C | | | | |
| HDX time course (min) | 0.5, 1, 2, 5 min | | | | |
| HDX control samples | We use the highest exchanging peptides as maximally deuterated controls | | | | |
| Back-exchange (mean / IQR) | 22%/8% | 22%/8% | 22%/8% | 22%/8% | 22%/8% |
| # of Peptides | 82 (RelA)<br>63 (p50) | 82 (RelA)<br>63 (p50) | 82 (RelA)<br>63 (p50) | 82 (RelA)<br>63 (p50) | 82 (RelA)<br>63 (p50) |
| Sequence coverage | 73.1% (RelA)<br>96.2% (p50) | 73.1% (RelA)<br>96.2% (p50) | 73.1% (RelA)<br>96.2% (p50) | 73.1% (RelA)<br>96.2% (p50) | 73.1% (RelA)<br>96.2% (p50) |
| Average peptide length / Redundancy | 13.5/2.86 (RelA)<br>14.1/2.89 (p50) | 13.5/2.86 (RelA)<br>14.1/2.89 (p50) | 13.5/2.86 (RelA)<br>14.1/2.89 (p50) | 13.5/2.86 (RelA)<br>14.1/2.89 (p50) | 13.5/2.86 (RelA)<br>14.1/2.89 (p50) |
| Replicates (biological or technical) | 3 technical replicates each | | | | |
| Repeatability (average standard deviation) | 0.060 (RelA)<br>0.060 (p50) | 0.060 (RelA)<br>0.060 (p50) | 0.060 (RelA)<br>0.060 (p50) | 0.060 (RelA)<br>0.060 (p50) | 0.060 (RelA)<br>0.060 (p50) |
| Significant differences in HDX (delta HDX > X D) | 0.5 D (99%CI) | | | | |

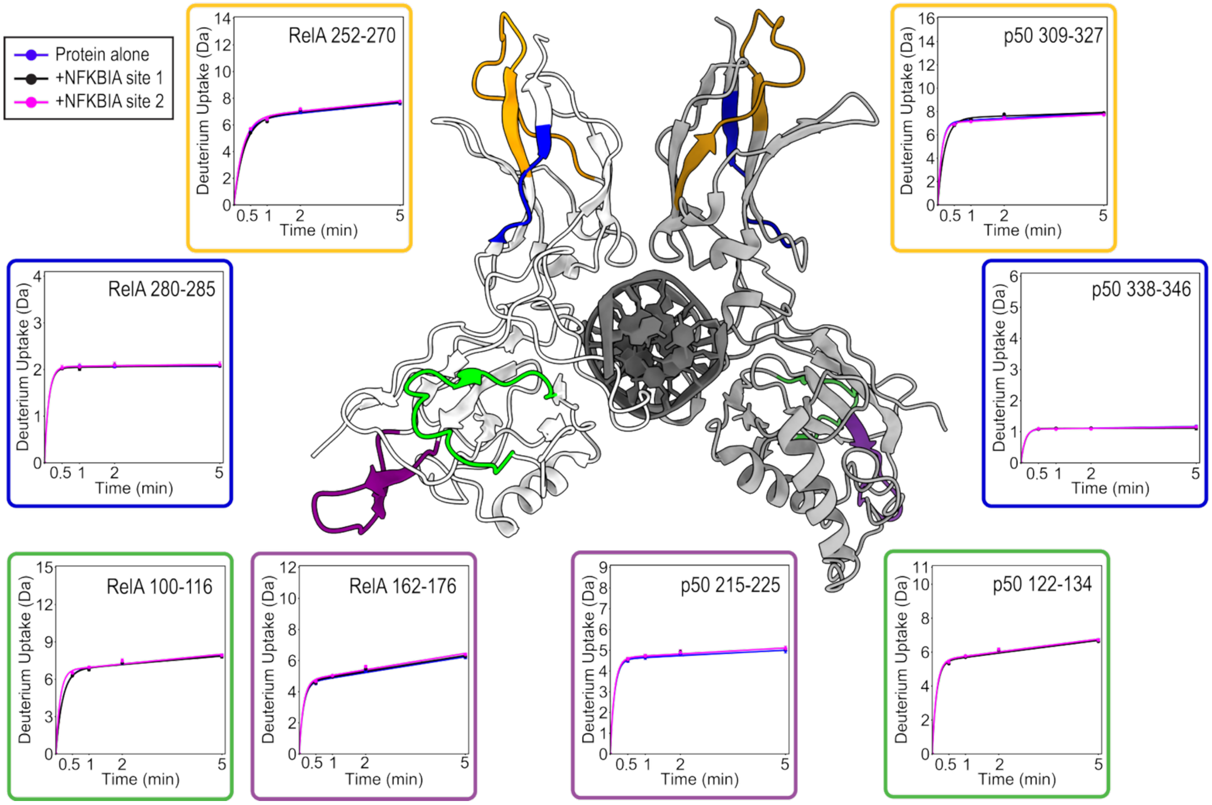


**Supporting Figure 1. Using HDX-MS, exterior regions of RelA/p50 alone vs. bound to NFKBIA site 1 and site 2 had unchanged structural dynamics outside of the DNA binding pocket.** HDX-MS was used to compare structural dynamics for regions of RelA/p50 outside of the DNA binding pocket. Using PDB 1LE5, regions of RelA (left, light gray) and p50 (right, dark gray) are color-coordinated to uptake plot outlines. Deuterium uptake plots for the region of free (blue), site 1-bound (black), or site 2-bound (pink) RelA/p50, with the subunit identity and residue range numbered in the top right of each plot. The representative regions of RelA/p50 highlighted here had no significant differences in deuterium uptake whether DNA-bound or free. The RelA transcription activation domain (not pictured) similarly showed no significant changes with DNA binding.

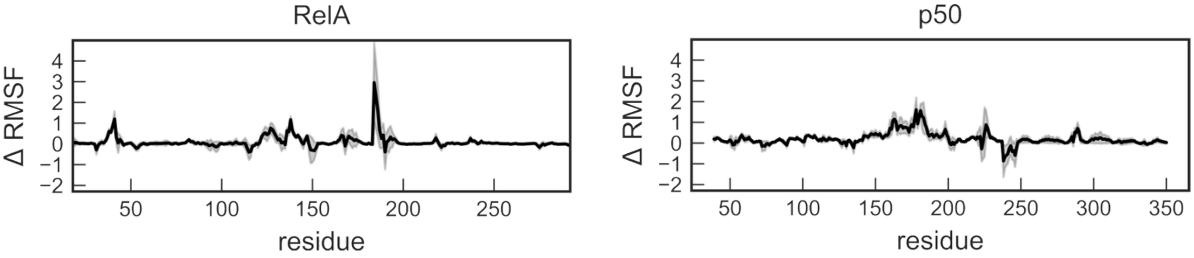


**Supporting Figure 2. RMSF differences between site 1- and site 2A-bound RelA/p50 residues indicate greater flexibility on half-site DNA.** Changes in the RMSF of RelA and p50 residues when bound to NFKBIA site 2A vs. site 1 $(\mathrm{RMSF}_{\mathrm{site\ 2A}} - \mathrm{RMSF}_{\mathrm{site\ 1}} = \Delta\mathrm{RMSF})$ were calculated from unweighted simulations. The standard error is shown as a gray envelope. Differences were primarily observed through the NTD and linker regions of RelA and p50, and were minimal through the dimerization domains (residues 195-291 and 249-350 of RelA and p50, respectively).

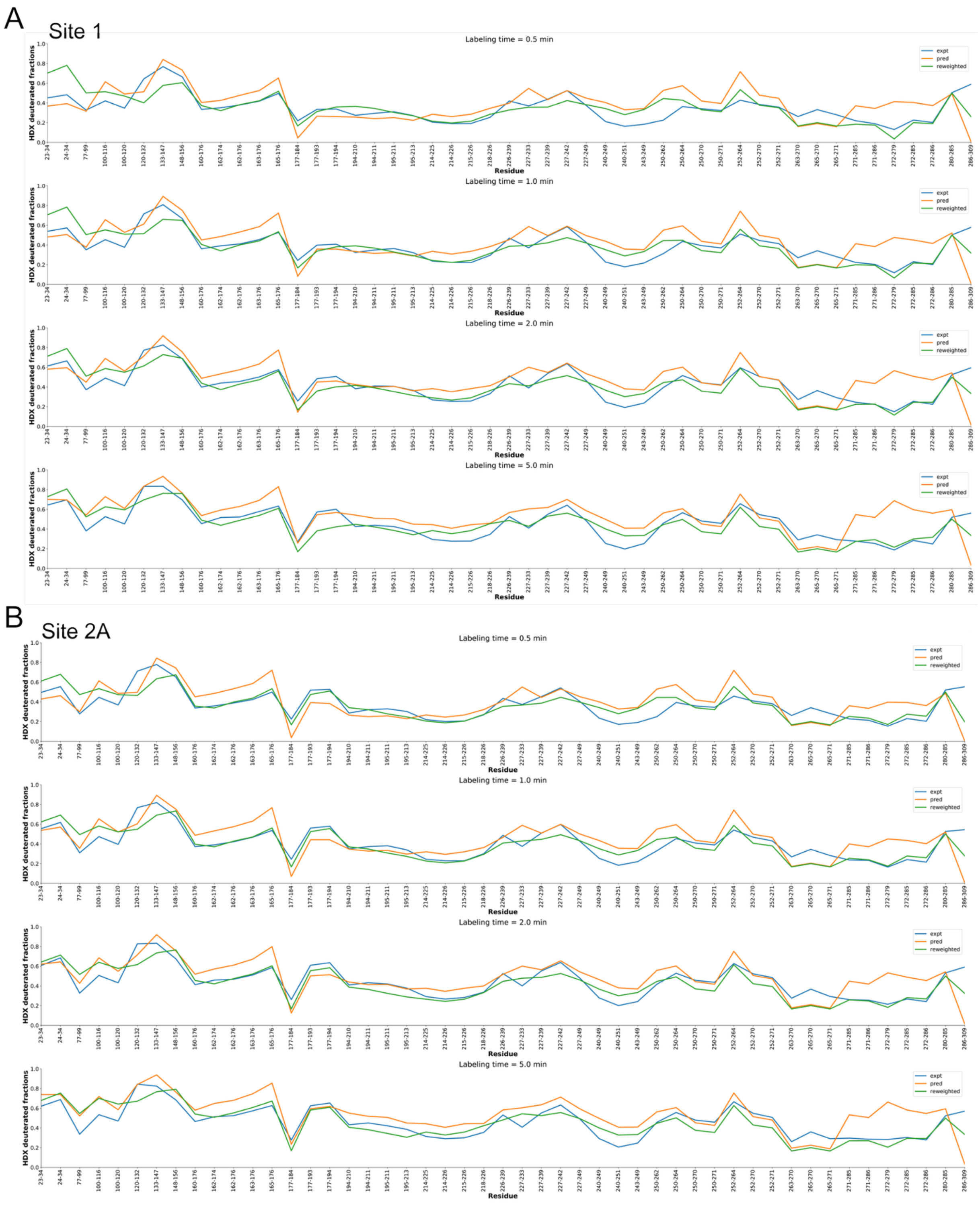


**Supporting Figure 3. For RelA, HDXer-predicted deuterium uptake matches well with experimental HDX-MS.** A deuterated fraction value was calculated from experimental HDX-MS data (blue) for each RelA peptide using actual deuterium uptake over maximum theoretical uptake. For the same peptides, deuterated fractions were predicted based on simulations (orange), and predicted again following reweighting (green) for site 1 (A) and site 2A (B).

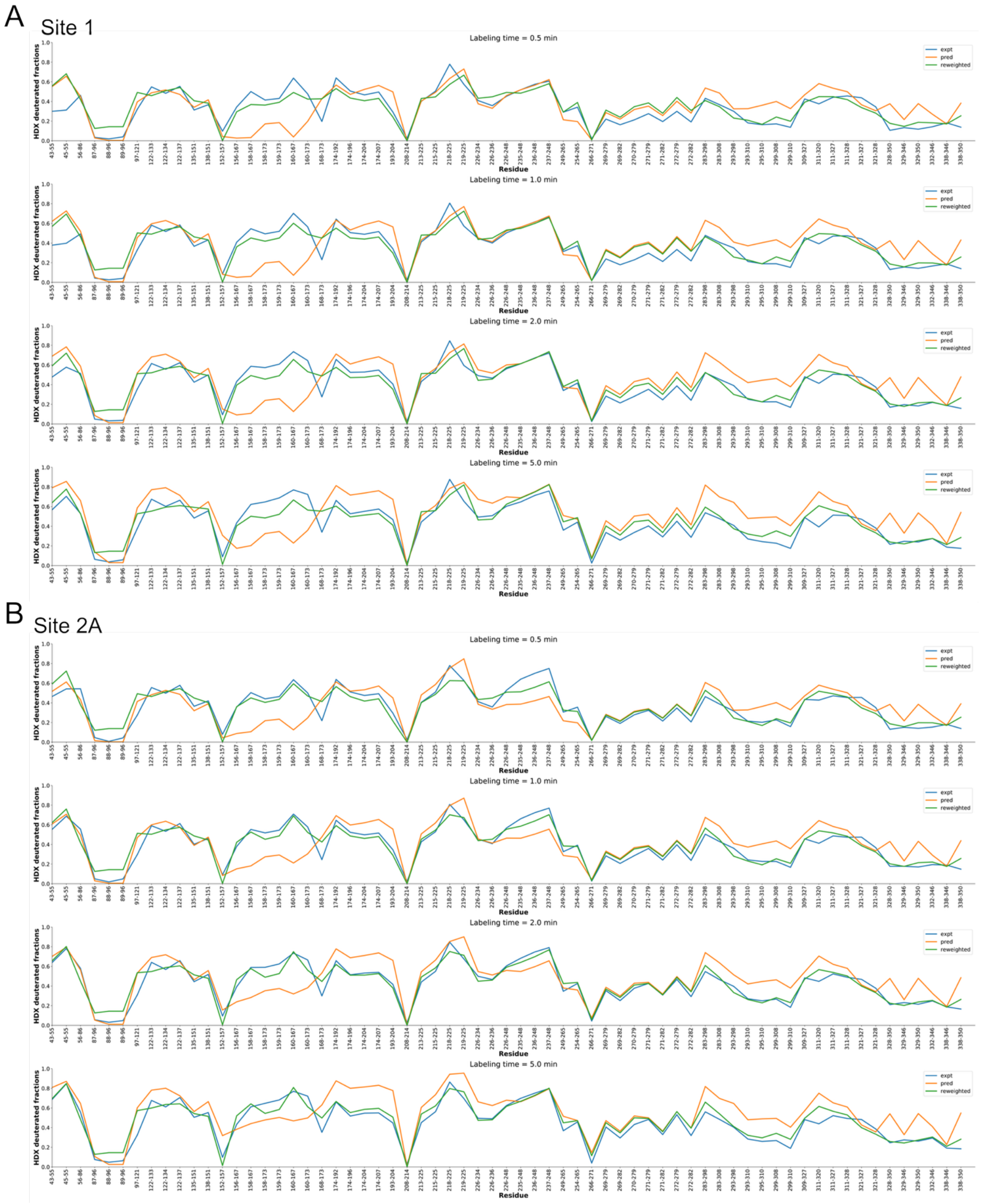


**Supporting Figure 4. For p50, HDXer-predicted deuterium uptake matches well with experimental HDX-MS.** A deuterated fraction value was calculated from experimental HDX-MS data (blue) for each p50 peptide using actual deuterium uptake over maximum theoretical uptake. For the same peptides, deuterated fractions were predicted based on simulations (orange), and predicted again following reweighting (green) for site 1 (A) and site 2A (B).

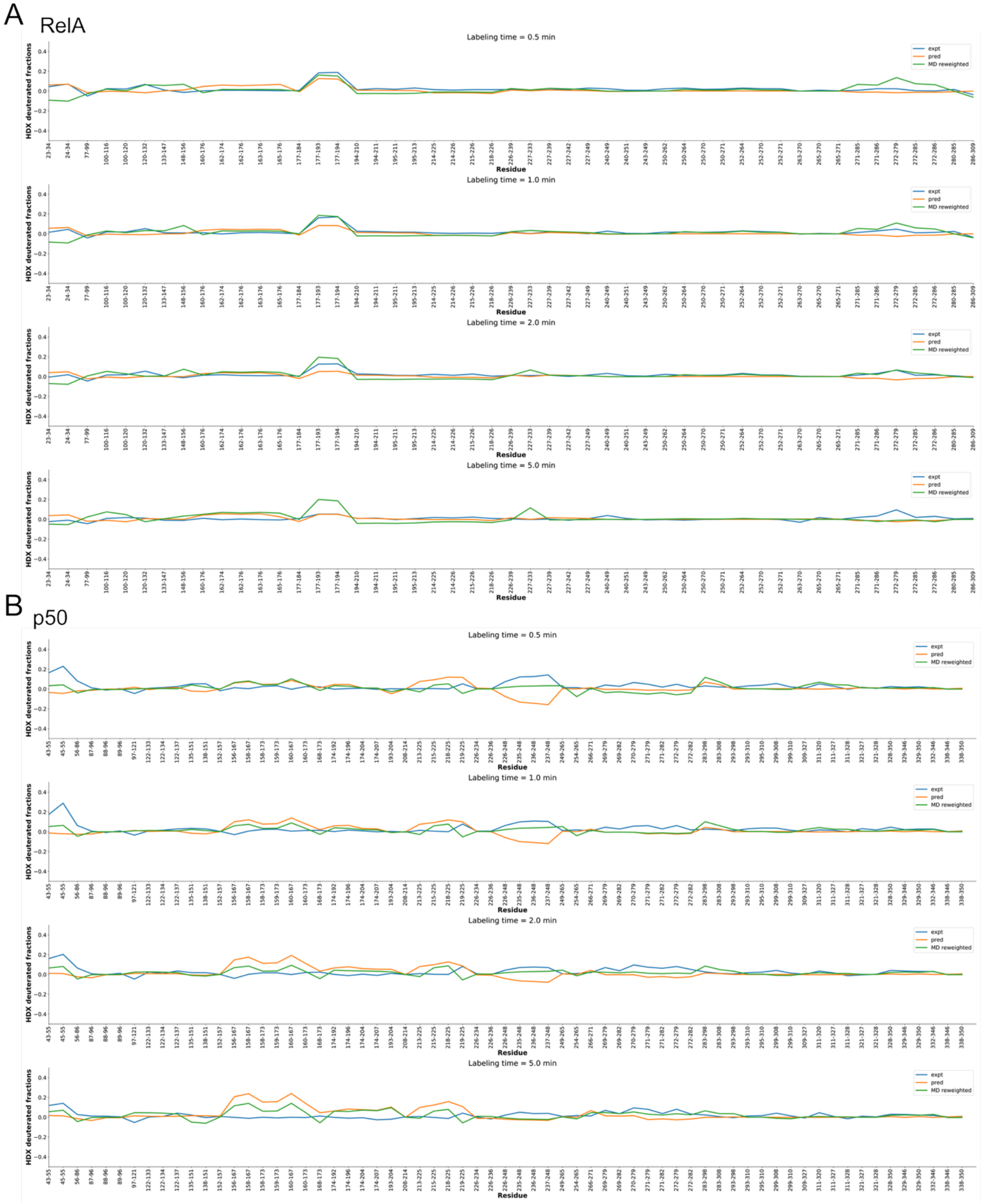


**Supporting Figure 5. HDXer-predicted differences in deuterated fraction between site 2A- and site 1-bound RelA/p50 match closely with experimental HDX-MS for RelA but fail to recapitulate results for the p50 linker.** A deuterated fraction difference value was calculated by subtracting values from Supporting Figure 3A from Supporting Figure 3B for RelA, and Supporting Figure 4A from Supporting Figure 4B for p50 ($\Delta \text{deuterated fraction} =$

$\text{deuterated fraction}_{site\ 2A} - \text{deuterated fraction}_{site\ 1}$). (A) Experimentally determined differences (blue) match closely with predicted differences (orange) and reweighted predictions (green) for RelA. (B) Experimental results (blue) do not match predicted differences (orange) through the p50 linker region (region 237-248), although the discrepancy is minimized upon reweighting (green).
